# Two distinct functional axes of positive feedback-enforced PRC2 recruitment in mouse embryonic stem cells

**DOI:** 10.1101/669960

**Authors:** Matteo Perino, Guido van Mierlo, Sandra M.T. Wardle, Hendrik Marks, Gert Jan C. Veenstra

## Abstract

Polycomb Repressive Complex 2 (PRC2) plays an essential role in development by catalysing trimethylation of histone H3 lysine 27 (H3K27me3), resulting in gene repression. PRC2 consists of two sub-complexes, PRC2.1 and PRC2.2, in which the PRC2 core associates with distinct ancillary subunits such as MTF2 and JARID2, respectively. Both MTF2, present in PRC2.1, and JARID2, present in PRC2.2, play a role in core PRC2 recruitment to target genes in mouse embryonic stem cells (mESCs). However, it remains unclear how these distinct sub-complexes cooperate to establish Polycomb domains. Here, we combine a range of Polycomb mutant mESCs with chemical inhibition of PRC2 catalytic activity, to systematically dissect their relative contributions to PRC2 binding to target loci. We find that PRC2.1 and PRC2.2 mediate two distinct paths for recruitment, with mutually reinforced binding. Part of the cross-talk between PRC2.1 and PRC2.2 occurs via their catalytic product H3K27me3, which is bound by the PRC2 core-subunit EED, thereby mediating a positive feedback. Strikingly, removal of either JARID2 or H3K27me3 only has a minor effect on PRC2 recruitment, whereas their combined ablation largely attenuates PRC2 recruitment. This strongly suggests an unexpected redundancy between JARID2 and EED-H3K27me3-mediated recruitment of PRC2. Furthermore, we demonstrate that all core PRC2 recruitment occurs through the combined action of MTF2-mediated recruitment of PRC2.1 to DNA and PRC1-mediated recruitment of JARID2-containing PRC2.2. Both axes of binding are supported by EED-H3K27me3 positive feedback, but to a different degree. Finally, we provide evidence that PRC1 and PRC2 mutually reinforce reciprocal binding. Together, these data disentangle the interdependent and cooperative interactions between Polycomb complexes that are important to establish Polycomb repression at target sites.

**Highlights:**

- Systematic analysis of Polycomb complex binding to target loci in mESCs using null mutations and chemical inhibition.
- PRC1, PRC2.1 and PRC2.2 are all mutually dependent for binding to chromatin, mediated in part by H3K27me3.
- PRC2.1 recruitment is dependent on MTF2
- PRC2.2 recruitment by JARID2 is dependent on PRC1 and largely redundant with recruitment by H3K27me3

## Introduction

Cell fate specification during embryonic development requires tightly controlled epigenetic programs. A key component safeguarding these processes is Polycomb Repressive Complex 2 (PRC2), an enzymatic protein complex that catalyses mono-, di- and trimethylation of histone 3 lysine 27 (H3K27me1/2/3), which plays an essential role in the establishment of cellular identity by ensuring proper gene silencing (Pengelly et al., 2013). The critical role of PRC2 during developmental processes is underscored by the embryonic lethality observed in mice lacking a functional PRC2 complex (Faust et al., 1998; O’Carroll et al., 2001; Pasini et al., 2007). PRC2 consists of the core subunits EED, SUZ12 and EZH2, the latter being the catalytic subunit. In addition, PRC2 contains multiple ancillary subunits exerting functions such as guiding PRC2 to target genes and modulating its enzymatic activity. These include PCL proteins (PHF1, MTF2 or PHF19), EPOP (also known as C17ORF96) and PALI1/2 (also known as C10ORF12), which together with the core subunits comprise PRC2.1. The core subunits can alternatively associate with JARID2 and AEBP2 in another PRC2 subcomplex, referred to as PRC2.2 (Conway et al., 2018; van Mierlo et al., 2019a).

Within mouse embryonic stem cells (mESCs), the PRC2 core complex is mainly associated with MTF2 and EPOP (PRC2.1), or with AEBP2 and JARID2 (PRC2.2) (Kloet et al., 2016). Alternative PRC2.1 complexes containing either PHF1 or PHF19, and/or PALI1/2 are less abundant. In recent years, our understanding of Polycomb regulation in terms of recruitment and enzymatic activity has significantly increased. First, it has been shown that PRC2 can be recruited by the facultative subunits MTF2 and JARID2 in mESCs, while ablation of either EPOP or AEBP2 does not affect PRC2 localization (Beringer et al., 2016; Casanova et al., 2011; Grijzenhout et al., 2016; Landeira et al., 2010; Li et al., 2017; Liefke et al., 2016; Son et al., 2013). Second, after the first establishment of PRC2 binding, the complex can self-reinforce and spread from its target sites through an allosteric positive feedback loop by binding of the EED WD40 domain to H3K27me3 (Margueron et al., 2009; Poepsel et al., 2018). As this mechanism is not sufficient for H3K27me3 maintenance during cell division (Laprell et al., 2017), this indicates the importance of continuous *de novo* recruitment of core PRC2 by its auxiliary subunits. Third, PRC2 can be recruited through non-canonical PRC1, which binds to non-methylated DNA via its subunit KDM2B, and catalyses ubiquitination of H2A (H2AK119ub). This mark, in turn, can be bound by JARID2, resulting in PRC2 recruitment (Cooper et al., 2016; Kalb et al., 2014; Tavares et al., 2012). Finally, the H3K27me3 mark can be bound by canonical PRC1 via CBX subunits, which may contribute to gene repression by chromatin compaction (Isono et al., 2013; Lau et al., 2017; Morey et al., 2012). The bulk of H2A ubiquitination, however, is mediated by variant PRC1 complexes that contain one of several PCGF proteins (Fursova et al., 2019).

It has become clear that MTF2 and JARID2 together are required for PRC2 recruitment to target genes in mESCs, as combined ablation of MTF2 and JARID2 in mESCs lack PRC2 recruitment to target genes (Oksuz et al., 2018). This seems to depend to a large extent on MTF2-mediated DNA binding with a relatively minor contribution of JARID2 (Casanova et al., 2011; Li et al., 2017; Perino et al., 2018). Yet, while MTF2 and JARID2 are mutually exclusive within PRC2 complexes, the absence of either of the two partially reduces the binding of the other (Perino et al., 2018). This suggests that PRC2.1 and PRC2.2 could directly or indirectly synergize in establishing Polycomb at target genes. Whether such a cooperativity exists, how it would materialize, what the relative contribution is of PRC2.1 and PRC2.2, and how PRC1 plays a role in this process remains to be defined. Here, we combine a range of Polycomb mutant ESCs with chemical inhibition of PRC1 and PRC2 to address the complex interactions of the Polycomb system using ChIP-sequencing. We assess the individual contributions of primary recruitment mechanisms established by JARID2, MTF2 and H3K27me3. Our data strongly indicates that PRC2.1 and PRC2.2 act synergistically for PRC2 recruitment, an interaction that is partially mediated through H3K27me3. Furthermore, our data indicate that H3K27me3-mediated recruitment of PRC2 can be compensated for by JARID2-mediated recruitment and vice versa. Moreover, we provide evidence that this apparent redundancy is mediated through JARID2 and PRC1-deposited H2AK119ub. Together, our data support a model in which core PRC2 recruitment requires the concerted action of MTF2 and JARID2, as well as EED binding to H3K27me3. These modes of recruitment can be subdivided into two major axes, of which one relies more on MTF2-mediated DNA binding, and the other to a larger extent on JARID2-H3K27me3-PRC1 mediated recruitment. Moreover, these different recruitment axes appear to carry different weights across the genome. The data presented here demonstrate that the interactions between PRC2 sub-complexes are tuned depending on the genomic region and highlight their relevance in establishing PRC2 binding at target sites.

## Methods

### Embryonic stem cell culture

Wildtype E14 ESCs (129/Ola background) and knockout ESCs were maintained in Dulbecco’s Modified Eagle Medium (DMEM) containing 15% fetal bovine serum, 10 mM Sodium Pyruvate (Gibco), 5 μM beta mercaptoethanol (BME; Sigma) and Leukemia inhibitory factor (LIF: 1000U/ml; Millipore). *Eed^-/-^* ESCs have been described by Schoeftner et al.(Schoeftner et al., 2006), *Jarid2^-/-^* ESCs have been described in Landeira et al. (Landeira et al., 2010), *Mtf2* knockout (*Mtf2^GT/GT^*) (Li et al., 2011) and *Ring1a^-/-^/ Ring1b^+/-^* ESCs (Endoh et al., 2008) were a kind gift from Haruhiko Koseki. *Ring1b* ESCs are knockout for *Ring1a* and trans-heterozygous for *Ring1b* (null/floxed). Full knockout of *Ring1b* was induced through treatment with Tamoxifen (OHT) for 2 days. To inhibit EED function, ESCs were treated with 10 μM EED226 (Qi et al., 2017) for 4 days. Complete removal of H3K27me3 was validated using western blot. To deplete H2AK119ub, mESCs were treated with 10 μM MG132 for 6 hours (Tavares et al., 2012). Complete removal of H2AK119ub was validated using western blot.

### Western blot and antibodies

Cell pellets were dissolved in RIPA buffer at a density of 10^4^ cells per μl and briefly sonicated to ensure proper cell lysis. Proteins denatured in SDS-PAGE gels were transferred onto PVDF membranes. Primary antibodies used were rabbit anti-MTF2 (ProteinTech; 16208-1-AP), rabbit anti-JARID2 (Novus Bio; NB100-2214), rabbit anti-H3K27me3 (Millipore; 07-449), rabbit anti-H3 (Abcam;1791). Secondary antibodies were HRP-conjugated anti-rabbit (Dako; P0217) and anti-mouse (Dako; P0161). Protein bands were visualized using Pierce ECL western blotting substrate (Thermo). Images were analysed using ImageJ.

### ChIP-sequencing

Cells were crosslinked in 1% PFA at room temperature for 10 min. The crosslinking reaction was halted using 1.25M glycine and cells were harvested by scraping in buffer B (0.25% Triton X-100, 10 mM EDTA, 0.5 mM EGTA, 20 mM HEPES). The suspension was centrifuged for 5 min at 1600 rpm, 4 °C and the pellet was resuspended in 30 ml buffer C (150 mM, 1 mM EDTA, 0.5 mM EGTA, 50 mM HEPES) and rotated for 10 min at 4 °C. The nuclei were centrifuged 5 min at 1600 rpm, 4 °C and resuspended in incubation buffer (0.15% SDS, 1% Triton X-100, 150 mM NaCl, 1 mM EDTA, 0.5 mM EGTA, 20 mM HEPES) supplemented with Protease inhibitor. Nuclei were sonicated using a Biorupter Pico to obtain chromatin with an enriched DNA length of 300 bp. The chromatin was snap frozen and stored at −80 °C until further use. For ChIP, sonicated chromatin was incubated overnight with the required antibody and pulled down using protein A/G magnetic beads (Perino et al., 2018). After washes, eluted chromatin was de-crosslinked overnight and purified with MinElute PCR Purification columns (Qiagen). After qPCR quality check for target enrichment, up to 5 ng/sample of ChIP was prepared for sequencing using the Kapa Hyper-prep Kit (Kapa Biosystems) using NEXTflex adapters (Bio Scientific), followed by 8-11 cycles amplification by PCR. After size-selection using E-gel (Invitrogen) to enrich for 300bp fragments, libraries were sequenced paired-end on an Illumina NextSeq500. qPCR analysis of ChIP DNA was performed with iQ SYBR Green Supermix (Bio-Rad) on a CFX96 Real-Time System C1000 Thermal Cycler (Bio-Rad). All the ChIP-Seq experiments in this study were performed at least in duplicate, from independent chromatin preparations.

### ChIP antibodies

ChIP was performed using 3 μl/sample of the following antibodies: MTF2 (Aviva System Biology ARP34292, lot QC49692-42166), H3K27me3 (Millipore 07-449, lot 2717675), EZH2 (Diagenode C15410039, lot 003), JARID2 (Novus Biologicals NB100-2214, Lot E2), RING1B (Abcam, AB3832 lot GR86503-25).

### Bioinformatic analysis

Data from Perino *et al.*,(Perino et al., 2018) were reprocessed in parallel with those of this study. To ensure maximum comparability (75bp single-end vs 42bp paired-end) and accurate quantification, all fastq files were trimmed to 42bp using fastx_trimmer (version 0.0.13.2), and in case of paired-end sequencing only read_1 was used analysis. All fastq files were mapped using bwa (version 0.7.10-r789), filtered to retain only uniquely mapping reads using mapping quality of 30 and samtools (version 1.7, flag -F 1024), then normalized for sequencing depth to produce bigwig. Peaks were called with MACS2-2.7 (Zhang et al., 2008) with qvalue 0.0001 using --call-summits for transcription factors and --broad for H3K27me3. Only peaks independently called in both replicates were used for downstream analysis. High-confidence peaks for each mark were obtained by merging peaks called in both replicates and overlapping by at least 50% of their length, and combined to obtain the list of all PRC2 peaks. Heatmaps of ChIP-Seq signal were generated using fluff v3.0.2 (Georgiou and van Heeringen, 2016), and clustered for dynamics using the “–g” option. ChIP metaplots were obtained with deeptools v 3.1.3 (Ramírez et al., 2016). Anatomy term enrichment was calculated using MouseMine (Motenko et al., 2015). RPKM bootstrapping analysis was performed using scipy (v 1.1.0). RPKM from the two independent ChIPseq replicates were combined into a single pool. Values were drawn from this pool, recorded, and returned, such that every value could be drawn multiple times. For each bootstrapping round a number of values matching the total number of PRC2 peaks was drawn, and the median plotted as one dot in the swarmplot. Confidence intervals (99.9%) were calculated from 100 bootstrapping events.

### Whole cell proteomes

Cell pellets were dissolved in RIPA buffer at a density of 10^4^ cells per μl and briefly sonicated to ensure proper cell lysis (van Mierlo et al., 2019b). Protein extracts (10 μg) were processed using Filter Aided Sample Preparation (FASP) and digested overnight with Trypsin. Peptide mixtures were desalted prior to LC-MS analysis. Thermo RAW files were analysed using MaxQuant 1.5.1.0 with default settings and LFQ, IBAQ and match between runs enabled. In Perseus, contaminant and reverse hits were filtered out. WT, MTF2 knockout and JARID2 knockout ESCs were grouped. Only proteins that had an LFQ value in at least one of the conditions were maintained. Missing values were imputed using default settings in Perseus.

## Results

### PRC2 recruitment mainly depends on MTF2

Recent advances have pinpointed three main recruitment mechanisms of PRC2: 1) DNA-mediated recruitment via MTF2; 2) recruitment via JARID2; and 3) H3K27me3-mediated recruitment via EED (Fig. 1a) (Cooper et al., 2016; Li et al., 2017; Margueron et al., 2009; Oksuz et al., 2018; Pasini et al., 2010; Perino et al., 2018). Currently, it remains unclear how these mechanisms relate to each other or cooperate in establishing PRC2 binding at target genes. To investigate this, we first evaluated whether these mechanisms act at the same genomic sites by performing chromatin immunoprecipitation followed by massive parallel sequencing (ChIP-seq) using antibodies against endogenous EZH2, H3K27me3, MTF2 and JARID2. We performed peak calling for EZH2 and determined the occupancy of H3K27me3, MTF2 and JARID2 on these peak sites, which revealed a near-perfect overlap (Fig. 1b). The same scenario was evident for peaks called for H3K27me3 or MTF2 (Fig. S1a,b). By contrast, we observed a large number of sharp JARID2 peaks with little or no occupancy of the other PRC2 subunits (Fig. S1c; cluster 3). This could indicate that JARID2 exerts functions independent of the PRC2 complex, as previously suggested in *Drosophila* (Herz et al., 2012). However, as these sites do not overlap with other PRC2 subunits, they were excluded from consideration in this context and only the remaining clusters (Fig. S1d,e) were used for subsequent analysis.

**Figure 1.**
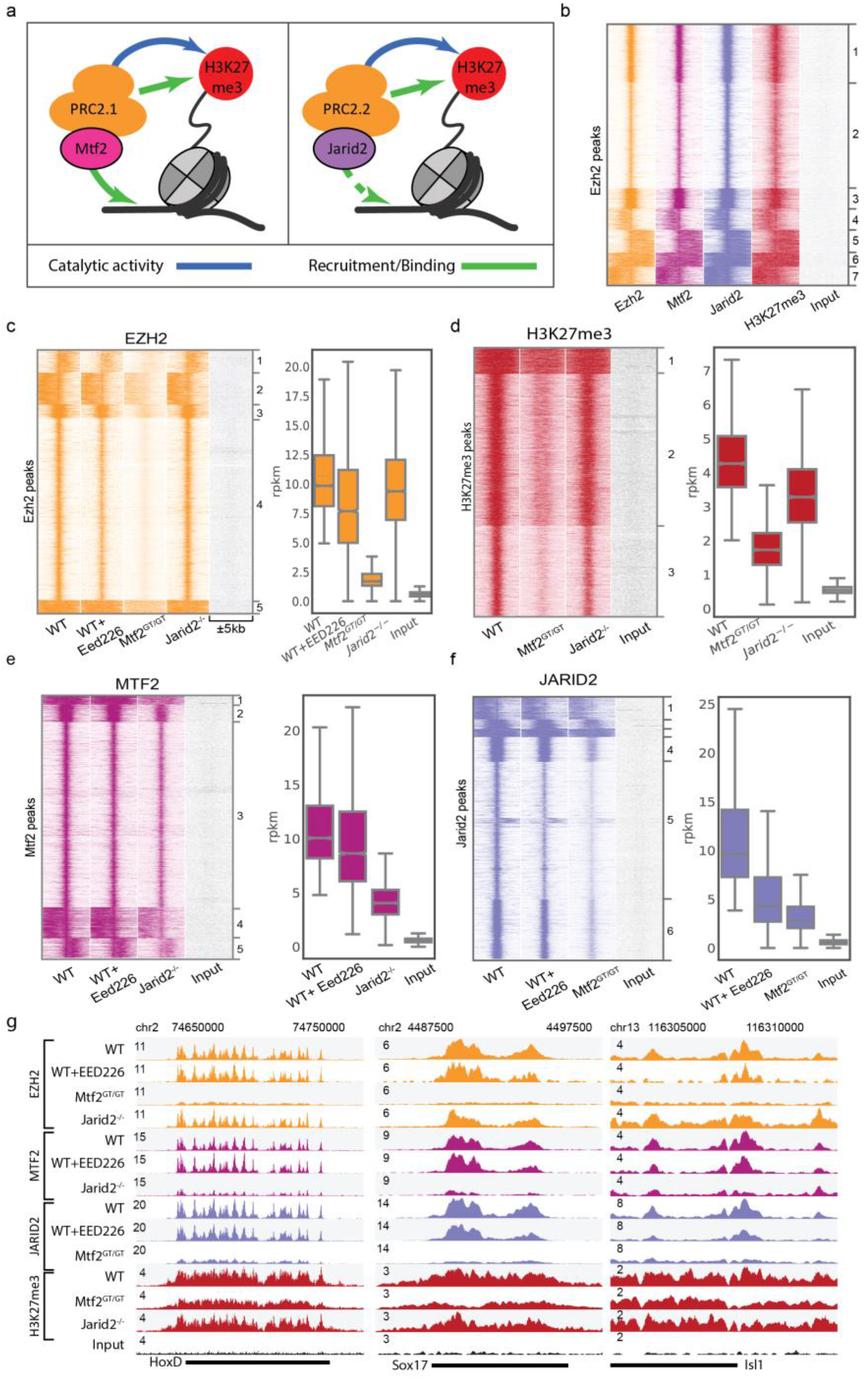
Canonical PRC2 recruitment largely relies on MTF2. **a)** Schematic representation of the recruitment of PRC2.1 and PRC2.2. MTF2 binds to DNA, while the EED subunit of core PRC2 (orange) binds to H3K27me3 as part of an allosteric feedback loop. The EZH2 subunit of core PRC2 catalyses H3K27 methylation. The PRC2.2 complex contains JARID2 but not MTF2. Both contain the core PRC2 subunits, however the interactions of the PRC2.1 and PRC2.2-specific subunits with chromatin are different. The arrow from JARID2 to DNA is dashed as DNA binding has been shown *in vitro* but not *in vivo* (Li et al., 2010) **b)** PRC2.1 (MTF2) and PRC2.2 (JARID2) co-localize to all EZH2 targets. **c-f)** Heatmap and rpkm quantification (boxplots) of PRC2 subunits and the catalytic product H3K27me3. EZH2 recruitment is heavily affected by the absence of MTF2, while JARID2 and H3K27me3 absence have minor effects (c). The effect of MTF2 and JARID2 on EZH2 recruitment is reflected on H3K27me3 deposition (d). MTF2 is marginally affected by H3K27me3 removal, but its binding is reduced to approximately half the WT level in the absence of JARID2 (e). JARID2 recruitment is strongly reduced in the absence of either H3K27me3 or MTF2 (f). ChIP profiles are highly reproducible (Fig S1h) **g)** Genome browser examples of PRC2 binding to classical Polycomb targets. Box plots represent median and interquartile range (IQR; whiskers, 1.5 IQR).

Next, we aimed to understand how MTF2, JARID2 and H3K27me3 are involved in recruitment of PRC2. First, we focused on MTF2 and JARID2 and used knockout mESCs for these subunits (*Mtf2^GT/GT^* and *Jarid2^-/-^*, respectively). These mESCs lack MTF2 or JARID2, respectively, but globally retain wildtype levels of core PRC2 subunits, as determined by quantitative mass spectrometry (Fig. S1f). ChIP-sequencing revealed a major reduction for EZH2 and H3K27me3 at target sites in *Mtf2* mutants, whereas the reduction in *Jarid2^-/-^* mESCs is minor (Fig. 1c,d). These observations are in line with previous reports attributing a prime role for MTF2 in PRC2 recruitment in mESCs (Li et al., 2017; Oksuz et al., 2018; Perino et al., 2018). To investigate whether PRC2.1 and PRC2.2 mediate recruitment of each other, we analysed the genomic locations bound by MTF2 and JARID2 in the knockout cells, which revealed that MTF2 and JARID2 mutually affect each other’s recruitment (Fig. 1e,f), with the most profound effect of *Mtf2* knockout on JARID2 recruitment (Fig. 1f). This suggests that the PRC2 sub-complexes directly or indirectly modulate their mutual recruitment. Next, to investigate the role of the allosteric EED feedback loop, we extended our analysis to wild type mESCs treated with the chemical inhibitor EED226. By binding the EED WD40 domain, EED226 hampers the binding of EED to H3K27me3 while simultaneously inducing a conformational change that impedes stimulation of the EZH2 catalytic activity by EED (Qi et al., 2017). Importantly, EED226 does not disturb physical associations between core PRC2 subunits (Qi et al., 2017). After confirming the complete absence of H3K27me3 in EED226-treated ES cells (Fig. S1g), we performed ChIP-seq for EZH2, MTF2 and JARID2. This revealed that in the absence of H3K27me3, EZH2 and MTF2 were retained on target sites at near wild type levels (Fig. 1c,e). JARID2 binding, instead, is >50% reduced by EED226 treatment (Fig. 1f), indicating that JARID2 recruitment to target sites partly relies on H3K27me3. Thus, the reduction of H3K27me3 in *Mtf2* mutant cells largely explains the reduction of JARID2 binding in this cell line, whereas the effect of JARID2 on MTF2 binding, which is largely independent of the levels of H3K27me3, might rely on a direct or indirect stabilization PRC2.1 association with chromatin. Examples of typical Polycomb targets are visualized in Fig. 1g. Taken together, these data confirm previous observations that primary recruitment of PRC2 occurs largely through MTF2 (Li et al., 2017; Perino et al., 2018), show its robustness in the absence of H3K27me3, and uncover a differential dependency on H3K27me3 levels for the PRC2.1 and PRC2.2 sub-complexes.

### Stratification of Polycomb binding sites reveals two major types of targets

We noticed that several of the clusters observed in Figure 1 showed distinct characteristics, such as the strength of binding or the width of the peaks (for example cluster 4 and 5 of the EZH2 ChIP-seq, Fig. 1c). Moreover, the consequences of removal of MTF2 or JARID2 appear different per cluster (c.f. the effect of MTF2 removal of cluster 2 and 4 in Fig. 1c, and cluster 3 and 5 in Fig. 1f, or the effect of JARID2 on clusters 2 and 5 in Fig. 1e). Furthermore, recent work implied that recruitment of MTF2 relies on the physical presence of EED only in a subset of PRC2 targets (Perino et al., 2018), which supports a hypothesis in which distinct modes of recruitment guide PRC2 to different genomic regions. To understand if and how PRC2 recruitment might be distinct depending on the genomic loci analysed, we included in our analysis MTF2 ChIP-seq data of mESCs lacking EED (Perino et al., 2018) (and therefore lacking the PRC2 core), BioCap data to identify regions free of DNA methylation (Long et al., 2013) (which is common for Polycomb targets and required for MTF2 binding to DNA (Perino et al., 2018)), and H3K4me3 ChIP-seq data of wild type ESCs (to identify bivalent promoter elements (Perino et al., 2018) that comprise the majority of Polycomb targets in mESCs (Brookes et al., 2012). We combined these data with those shown in Figure 1 and clustered them on the common set of PRC2-bound regions, revealing six major clusters (Fig. 2a). Clusters 1-4 display strong BioCap and H3K4me3 signals and are likely bivalent promoters (Bernstein et al., 2006), whereas clusters 5-6 show relatively low BioCap and H3K4me3 signals and could comprise silenced genes. We observed that the consequences of the perturbations for PRC2 recruitment varied per cluster (Fig. 2b, S2a). A notable example includes the H3K27me3 signal, which is affected more in clusters 1-4 (reduced to 6-27%) compared to cluster 5-6 (48-56%) in *Mtf2^GT/GT^* ESCs (Fig. 2b, top right). Similar patterns hold true for EZH2 (Fig. 2b, top left; 9-12% versus 23-26%) and JARID2 recruitment (Fig. 2b, bottom right; 11-19% versus 30-34%). In addition, removal of EED results in very strong reduction of MTF2 recruitment in all clusters. We recently found that MTF2 is recruited to CpGs in the context of a specific shape of the DNA, characterized by reduced helix twist (Perino et al., 2018). Therefore, we analysed the DNA shape of the genomic sequences in each cluster. This revealed that shape-matching GCG trinucleotides are much more prevalent in cluster 1-4 (Fig. 2c), which fits the higher dependence of MTF2 for PRC2 recruitment in these clusters (Perino et al., 2018). Together, these analyses indicate that cluster 1-4 rely relatively more on MTF2-mediated recruitment compared to cluster 5-6.

**Figure 2.**
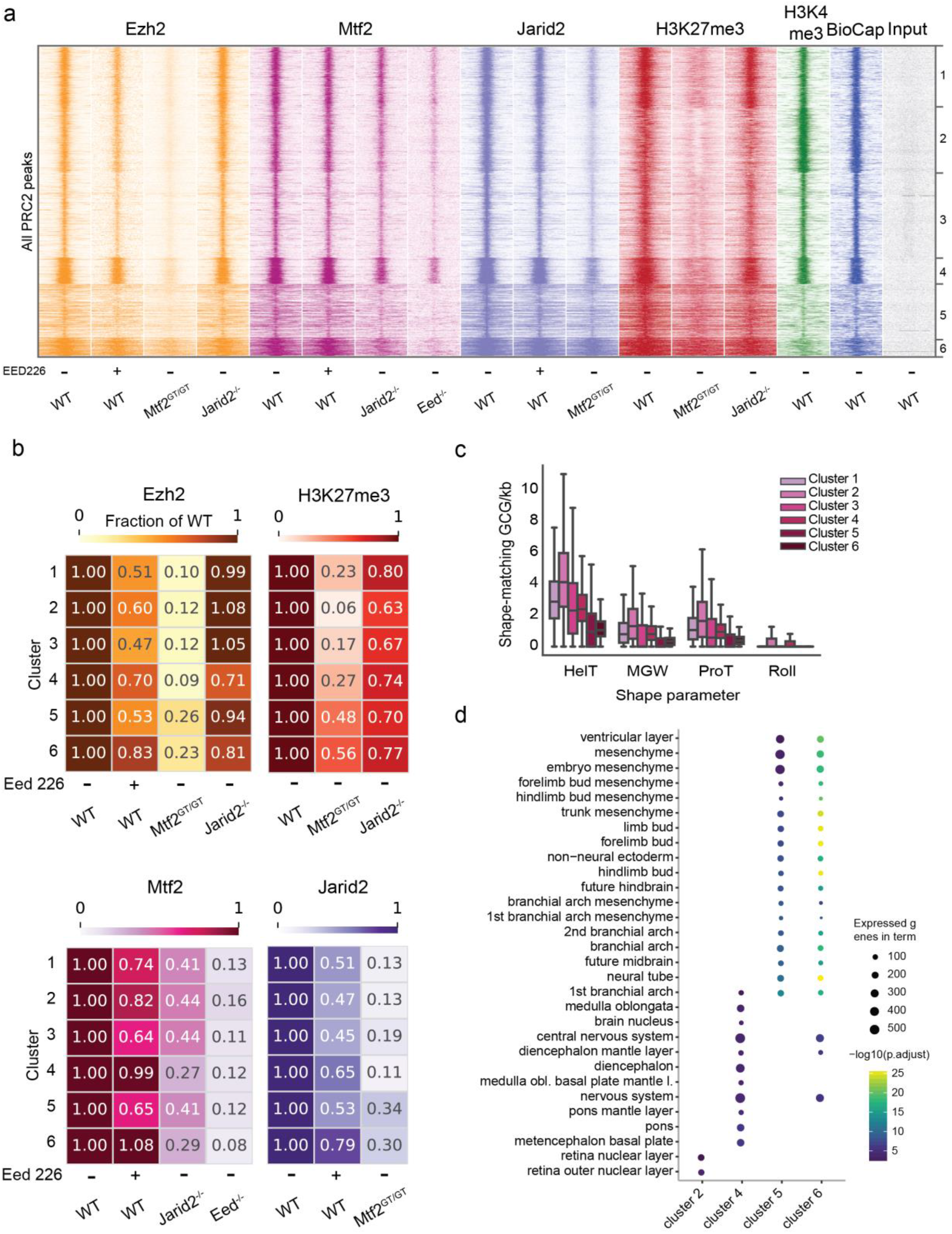
Identification of two distinct classes of Polycomb target regions, which rely on different mechanisms of PRC2 recruitment. **a)** Clustering of all PRC2 targets using ChIPseq data in multiple PRC2 mutants. Cluster 1-4 are unmethylated CpG islands (strong BioCap) signal, showing bivalent marks in WT (H3K4me3 and H3K27me3). These regions display heavy reduction of EZH2 recruitment in the MTF2 mutant, milder effects of H3K27me3 absence (EED226 treatment), and little or no effect of JARID2 absence. The intensity of MTF2 binding depends on both H3K27me3 and JARID2 but binding is still clearly detectable even in the absence of PRC2 core (*Eed^-/-^*) indicating a primary binding to DNA, reinforced by other mechanisms, such as JARID2-mediated recruitment, which in turn also depends on both H3K27me3 and MTF2. Cluster 5 and 6 have lower BioCap and H3K4me3 signal, and, while still affected by the absence of MTF2, this has a much less marked effect on recruitment of both EZH2 and JARID2, and on H3K27me3 deposition. **b)** WT-normalized, input-subtracted RPKM quantification of signal shown in (a). **c)** Quantification of GCG trinucleotides matching DNA shape requirement for MTF recruitments as defined in Perino et al, 2018. Cluster 1-4 are strongly enriched in shape-matching GCGs, indicating potential for strong DNA-mediated MTF2 recruitment. **d)** Enrichment of anatomical terms in the genes associated with peaks in the six clusters. Enrichment within PRC2 targets. Cluster 4 show strong enrichment for CNS structures, cluster 5 and 6 for limb and branchial arches tissues and mesenchyme. See Fig S2b for the full overview.

As previous reports suggested that Polycomb target sites contain distinct gene sets (Brookes et al., 2012), we tested whether cluster 1-4 and 5-6 also consisted of different sets of genes. When comparing with all the mouse genes, we observed that every cluster was enriched for genes associated with the development of body structures (Fig. S2b), as is characteristic for Polycomb genes in general (Brookes et al., 2012). Next, we stratified the clusters by calculating the enrichment among PRC2 targeted genes. We observed that cluster 5 and 6 are strongly enriched for genes related to body plan, limb, mesenchyme and branchial arches development (Fig. 2d), while cluster 2 and 4 show a stronger enrichment for neural structures (Fig. 2d). In addition, all Hox genes, which are considered highly conserved master regulators of embryonic development, are exclusively present in cluster 5-6. Collectively, these analyses identify two distinct classes of Polycomb target regions.

### JARID2 and H3K27me3 are redundant for PRC2 recruitment

Our analyses allowed us to investigate the individual contributions of MTF2, JARID2 and H3K27me3 for PRC2 recruitment. However, ablation of single subunits individually does not exclude the possibility of compensation by other factors. Thus, we combined knockouts of MTF2 and JARID2 with inhibition of H3K27 methylation. We treated *Mtf2^GT/GT^* ESCs with EED226 to remove H3K27me3, only leaving JARID2-mediated recruitment intact. Similarly, we combined removal of JARID2 with EED226 treatment, which leaves only the contribution of MTF2-mediated recruitment (cf. Fig 1a). In both situations, treatment with EED226 resulted in the complete removal of H3K27me3 (Fig S3). We examined the effect on core PRC2 recruitment to target genes by performing ChIP-sequencing of EZH2 in *Mtf2^GT/GT^*+EED226 ESCs and *Jarid2^-/-^*+EED226 mESCs. Inspection of the EZH2 signal revealed a relatively minor further decrease of EZH2 recruitment in *Mtf2^GT/GT^*+EED226 mESCs, compared to the already severe phenotype caused by MTF2 depletion alone (Fig 3a,b). Interestingly, although the absence of JARID2 alone had no effect (Fig. 3a,b; clusters 1-3, 5) or only a moderate effect (Fig. 3a,b; clusters 4, 6) on EZH2 recruitment, and the absence of H3K27me3 resulted in only a modest effect in all clusters, their combined ablation resulted in a dramatic decrease of EZH2 recruitment (Fig 3a,b). This could suggest that JARID2 and H3K27me3 are redundant for PRC2 recruitment or can compensate for each other. In addition, this demonstrates that MTF2-mediated recruitment by itself is not sufficient to establish full Polycomb recruitment, but requires PRC2.2 and/or the EED feedback loop. We extended our analyses by performing ChIP-sequencing for JARID2 in *Mtf2^GT/GT^*+EED226 mESCs and for MTF2 in *Jarid2^-/-^* +EED226 mESCs. Removal of both JARID2 and H3K27me3 further reduced MTF2 recruitment, and especially in cluster 5-6 MTF2 recruitment was near-zero (Fig 3c-e). This shows that these loci recruit MTF2 (PRC2.1) indirectly through PRC2.2 and the EED-positive feedback loop. This is in agreement with the observations that the loci in these clusters do not recruit MTF2 in the absence of PRC2 (*Eed^-/-^* mESCs) and do not show any enrichment for GCG trinucleotides with a DNA shape preferred by MTF2 (Fig 2a,d). When focusing on JARID2 in *Mtf2^GT/GT^*+EED226 ESCs, we observed a reduction of recruitment in all clusters although the decrease was marginally stronger in cluster 5-6 (Fig 3d). Together, these data uncover an important contribution of the EED-H3K27me3 interaction to PRC2 recruitment, in particular for PRC2.2, and show that the relative importance of PRC2.1 and PRC2.2 differs across the genome.

**Figure 3.**
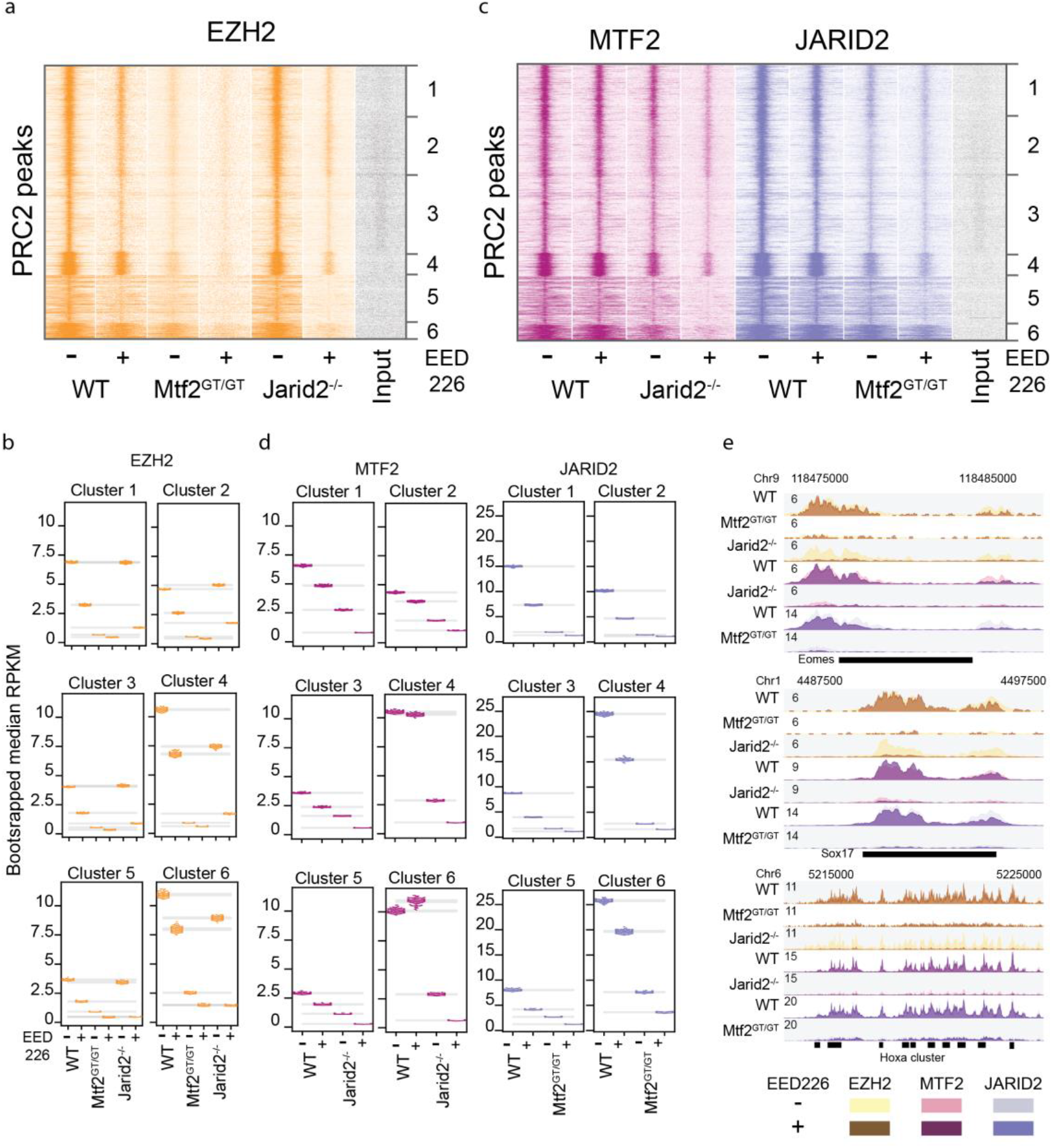
The H3K27me3 feedback loop and JARID2 are mutual backup for PRC2 recruitment. **a)** Heatmap showing the cluster specific effect of H3K27me3 depletion on the binding of EZH2. WT and MTF2^GT/GT^ show mild reduction of EZH2 binding when treated with EED226 inhibitor, while the treatment is highly synergistic with the depletion of JARID2. **b)** Bootstrapping-based RPKM quantification (methods) of the signal in (a). Each coloured dot represent the median of one round of bootstrapping, grey bar represent 99.9% confidence interval for the mean of bootstrapped values in each condition and cluster. **c)** Treatment with EED226 further affected MTF2 recruitment in *Jarid2^-/-^* and JARID2 recruitment in *Mtf2^GT/GT^*, with the former leading to recruitment patter closely resembling the *Eed^-/-^* line (cf. Fig2a), highlighting the recruitment differences between cluster 1-4 and 5-6. **d)** Bootstrapping-based RPKM quantification (methods) of the signal in (c) similar as in 3b. **e)** Genome browser view of example Polycomb targets. For each genotype two tracks are overlaid: the darker colour represent EED226 treated samples, the lighter colour untreated cells.

### JARID2 recruitment is largely dependent on PRC1

Our analyses indicate that the allosteric feedback loop mediated by EED seems to directly or indirectly buffer the absence of JARID2 and vice versa. We hypothesized this could be mediated through PRC1, as the catalytic subunit RING1B can deposit H2AK119ub which can in turn mediate JARID2 recruitment (Blackledge et al., 2014; Cooper et al., 2016). As PRC1-dependent PRC2 recruitment is mediated through H2AK119ub and JARID2, a scenario in which H3K27me3 and H2AK119ub are simultaneously absent might phenocopy the effect of *Jarid2^-/-^*+EED226 ESCs. To test this, we used *Ring1a/b* double mutant mESCs treated with EED226 (*Ring1a/b^-/-^*+EED226) and performed ChIP-sequencing of EZH2, MTF2 and JARID2 in these ESCs. Interestingly, we observed that the EZH2 and MTF2 profiles obtained in *Jarid2^-/-^*+EED226 and *Ring1a/b^-/-^*+EED226 were almost indistinguishable (Fig 4a-c, light and dark blue lines in Fig. 4b, Fig S4), and also JARID2 binding was affected in *Ring1a/b^-/-^*+EED226 cells (Fig 4d, Fig S4). Of note, while the residual JARID2 recruitment in *Ring1a/b^-/-^*+EED226 and *MTF2^GT/GT^*+EED226 is comparable for cluster 1-4, cluster 5-6 seems to depend more on PRC1 and on PRC2.2 (Fig 4d lower panels, Fig S4), further supporting a minor role to MTF2 for PRC2 recruitment at these locations. These observations strongly suggest that JARID2 and PRC1 act along same recruitment axis. As low levels of residual JARID2 recruitment is observed in clusters 1-4 in the *Ring1a/b*^-/-^+EED226 condition, additional mechanisms might recruit JARID2, for example binding of JARID2 to DNA (Li et al., 2010) or RNA (Brockdorff, 2013; Kaneko et al., 2014). To extend our analysis on PRC1-PRC2 interdependencies and disentangle the roles of H3K27me3 versus PRC2 subunits in PRC1 recruitment, we performed RING1B ChIP-seq in WT, *MTF2^GT/GT^*, and *Jarid2^-/-^* in the presence of EED226. Removal of H3K27me3 in wild type ESCs had only limited effect on RING1B recruitment (Fig 4e-g, Fig S5). However, the combined absence of H3K27me3 and either MTF2 or JARID2 results in strong reduction of RING1B binding (Fig 4e-g). While the Polycomb dogma posits that PRC1 and PRC2 do not physically interact and mutually affect each other only via their catalytic products (respectively H2AK119ub and H3K27me3), these data suggest that PRC2 also contributes to PRC1 recruitment independently of H3K27me3. For example, it is conceivable that the physical presence of PRC2 at target genes (which is strongly reduced in *Mtf2^GT/GT^*+EED226 and *Jarid2^-/-^*+EED226 ESCs) stabilizes PRC1 binding to chromatin by affecting chromatin compaction (Isono et al., 2013).

**Figure 4.**
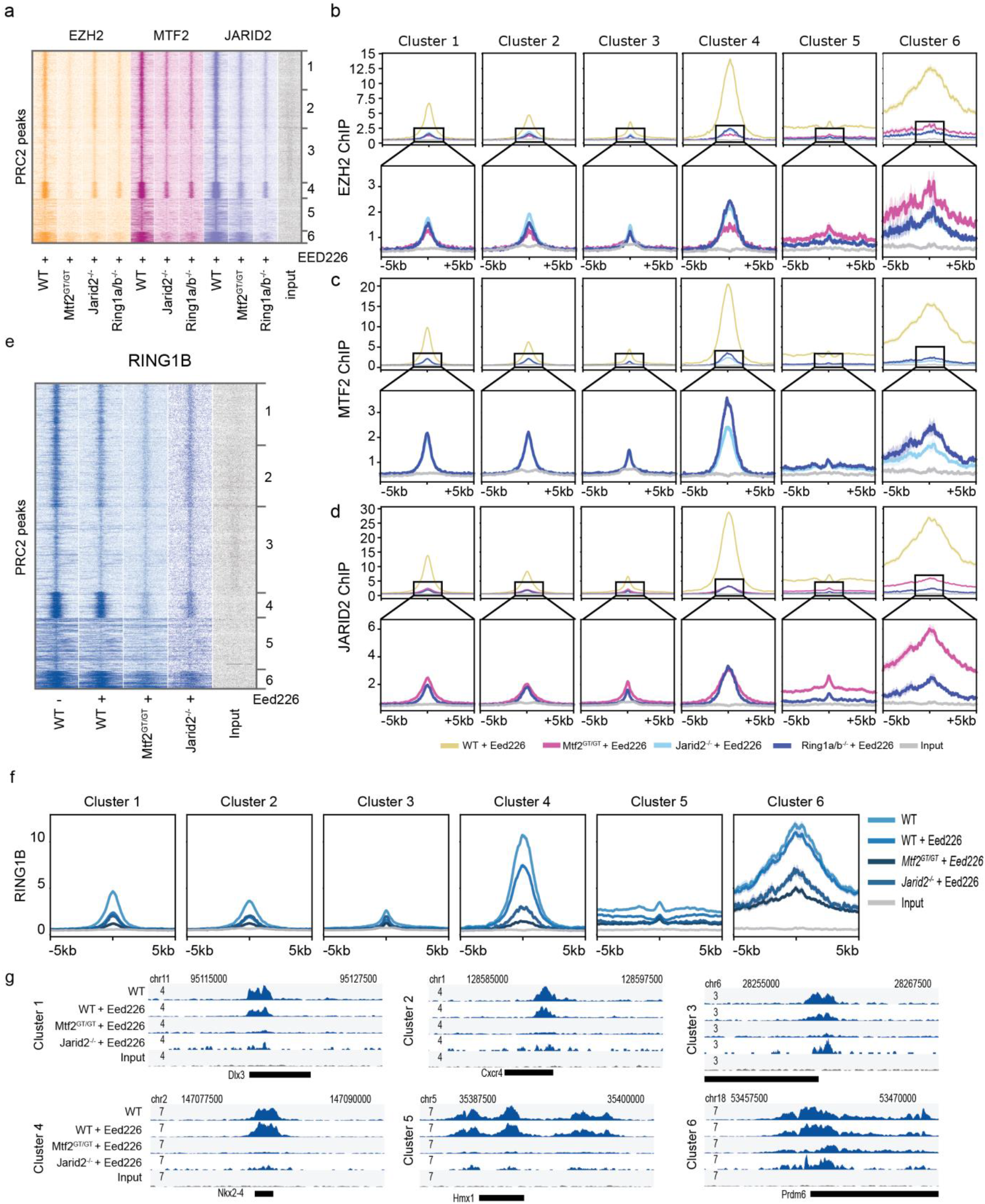
JARID2 recruitment is largely dependent on PRC1. **a)** Heatmap showing EZH2, MTF2 and JARID2 binding in the absence of H3K27me3 in PRC2 and PRC1 mutant lines. In the absence of H3K27me3, JARID2 and RING1A/B mutant phenocopy each other with regard to EZH2 and MTF2 binding, suggesting JARID2 and RING1B act in the same PRC2 recruitment mechanism. JARID2 recruitment is also strongly affected by the absence of RING1A/B, in line with the JARID2-mediated PRC2 recruitment via binding to PRC1-deposited H2AK119ub. **b-d)** Average plot of the ChIP signal shown in (a), for EZH2 (b) MTF2 (c) and JARID2 (d) centred on called peaks. Lower panels represent the same data with cropped y axis, for better visualization. **e)** Heatmap showing Ring1b binding in the discussed conditions. Ring1b is only mildly affected by removing H3K27me3 using EED226 (~40%). Binding is further attenuated in MTF2 and JARID2 mutant ESCs. **f)** Average plot of the ChIP signal shown in (e), centred on called peaks. **g)** Examples of loci of the data as shown in (e).

### PRC2 recruitment is mediated through a combined action of PRC2.1 and PRC2.2

Finally, we investigated whether the residual EZH2 recruitment observed in *Ring1a/b^-/-^*+EED226 and *Jarid2^-/-^*+EED226 mESCs was mediated through MTF2. To do so, we used *Mtf2^GT/GT^* ESCs in which we removed H3K27me3 using EED226 and additionally H2AK119ub using MG132 (Tavares et al., 2012) (*MTF2^GT/GT^*+d.i.; double inhibition), and performed ChIP-sequencing for EZH2. In this triple ablation condition, the recruitment of EZH2 to target genes was completely abrogated in all clusters (input levels, Fig 5, Fig S5). Collectively, these analyses demonstrate that the combined action of MTF2, the allosteric EED feedback loop and PRC1-mediated recruitment of JARID2-containing PRC2 sub-complexes are required for PRC2 recruitment in mESCs.

**Figure 5.**
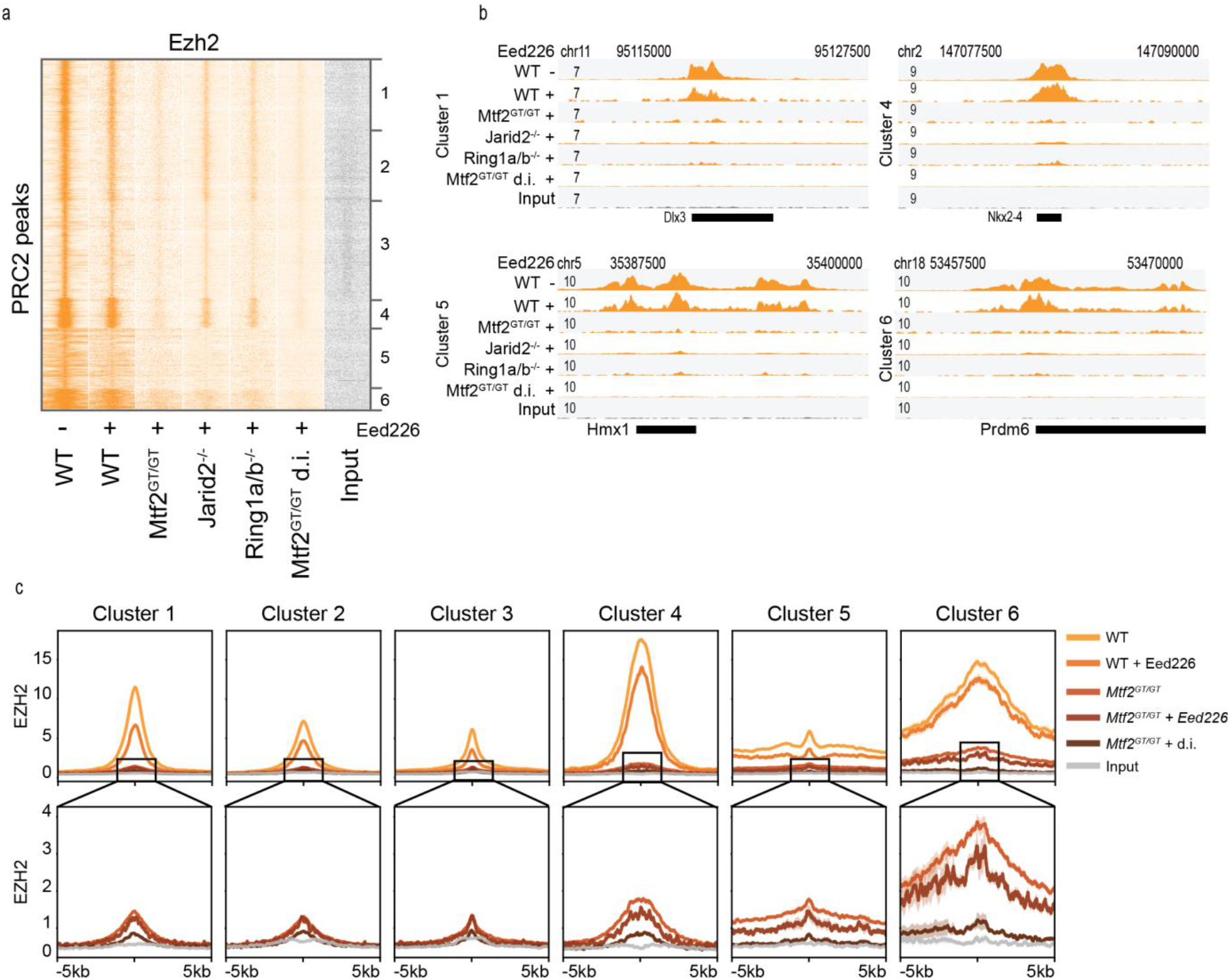
EZH2 recruitment depends on MTF2, H3K27me3 and H2AK119ub. **a)** Heatmap of ChIP-seq signal for EZH2 in multiple conditions including *Mtf2* cells with double inhibition (d.i.) using EED226 (to remove H3K27me3) and MG132 (to remove H2AK119ub). **b)** Example loci of the data shown in (a). **c)** Average profiles of the ChIP signal shown in (a), centred on called peaks. Lower panels represent the same data with cropped y axis, for better visualization.

## Discussion

The mechanisms that guide and maintain PRC2 at target sites have been the focus of extensive research, yet have long remained enigmatic. Although the allosteric feedback loop mediated by EED is important for spreading of PRC2 away from its initial nucleation site (Margueron et al., 2009), the mere presence of H3K27me3 is not sufficient to maintain PRC2 at its target genes (Laprell et al., 2017). This indicates that continuous DNA-mediated and Polycomb target-specific recruitment or stabilization is required to attract PRC2 to newly replicated chromatin fibres (Laprell et al., 2017). The recent discoveries of facultative PRC2 subunits and the presence of functionally distinct sub-complexes has greatly advanced our understanding of PRC2 recruitment and maintenance (Hauri et al., 2016; Smits et al., 2013). In particular, individual ablation of all prime facultative subunits in ESCs revealed a major role for MTF2 in PRC2 recruitment which, together with JARID2, mediates the initial PRC2 binding to the initiation sites (‘nucleation sites’) (Li et al., 2010, 2017; Oksuz et al., 2018; Perino et al., 2018).

In this study, we have dissected the various mediators of PRC2 recruitment. Our analyses confirm previous observations that MTF2 is required for a significant proportion of PRC2 recruitment(Li et al., 2017; Perino et al., 2018) and extend recent work highlighting that the remaining recruitment is mediated mostly via JARID2 (Oksuz et al., 2018). We show that MTF2 and JARID2 mutually modulate each other’s recruitment, partly through the EED feedback loop and in part through PRC1. Our work uncovers significant buffering and positive feedback in recruitment of PRC2. There are two main functional axes of primary PRC2 recruitment in mESCs, working through MTF2 and JARID2-PRC1, both of which are enforced by H3K27me3-EED positive feedback. The relative weight of these two mechanisms, however, depends on the genomic location (Fig 6). Polycomb target regions can indeed be sub-divided into (at least) two major categories. The largest group (in this study cluster 1-4 from Fig 2a onwards) contains mainly bivalent genes which rely more on PRC2.1-mediated recruitment. At these locations, a limited amount of MTF2 is sufficient to kick start recruitment, which is reinforced by EED feedback loop and PRC2.2, with partially redundant effects. Therefore, only combined ablation of both JARID2 and H3K27me3 dampens recruitment to the levels mediated by primary MTF2-dependent recruitment alone. Hence the simultaneous absence of MTF2, H3K27me3 and H2AK119ub is required to abolish all core PRC2 enrichment from target regions in mESCs. The smaller group (in this study cluster 5-6), instead, relies more on PRC1 and PRC2.2, and contains very lowly expressed and developmentally relevant genes such as all the Hox genes. Here, PRC1 activity is required to induce JARID2 and PRC2.2 recruitment, providing an alternative recruitment path to MTF2-dependent binding described above. MTF2 still binds to these locations, but does so indirectly, as shown by the loss of MTF2 in *Eed^-/-^*, *Jarid2^-/-^* +EED226 and *Ring1b^-/-^* +EED226, and supported by the sparse presence of DNA shape-permissive CG sequences, insufficient to achieve sustained DNA-driven MTF2 recruitment.

Beside the more intuitive effect of MTF2 on JARID2 recruitment, we also observed that JARID2 depletion affects MTF2 binding, but this is only partially mediated via H3K27me3, as shown by EED inhibition. Intriguingly, while affecting MTF2, the absence of JARID2 alone has a minimal effect on EZH2 recruitment. A potential explanation could be that hybrid PRC2.1/2.2 complexes containing AEBP2 and MTF2 form under these conditions, similarly to the JARID2-MTF2-containing hybrid complexes in AEBP2 mutant mESCs (Grijzenhout et al., 2016). The formation of hybrid complexes could sequester the complex or inhibit MTF2 recruitment.

The observations in the current study further substantiate previous work showing that the role of PRC1 and PRC2 are large intertwined, as both complexes can be recruited independently, but simultaneously modulate their mutual recruitment (Blackledge et al., 2014; Morey et al., 2013; Tavares et al., 2012). Our analyses of EED226-treated mESCs reveals that ~40% of PRC1 recruitment depends on the presence of H3K27me3 (Fig 4e-f), probably through canonical complexes containing CBX7 (Morey et al., 2012, 2013). The remainder of PRC2-independent PRC1 is likely recruited via KDM2B-mediated DNA binding, which is in line with previous observations showing that ~60% of RING1B recruitment is mediated by KDM2B (Farcas et al., 2012; Wu et al., 2013). Surprisingly, our analyses reveal that MTF2 and JARID2 deficient mESCs treated with EED226 show a more profound decrease (> ~40%) of RING1B occupancy at target genes. This could indicate either an as of yet unknown link between PRC1 and PRC2, or alternatively, a stabilization of KDM2B-mediated recruitment to DNA by the physical presence of core PRC2, that would increase the residence time of PRC1 on chromatin (Oksuz et al., 2018). Together, these observations further corroborate the hypothesis that PRC1 and PRC2 can bind autonomously, but are synergistic for their reciprocal recruitment.

Collectively, the observations here provide novel insights into Polycomb recruitment in ESCs and provide a model in which PRC2 recruitment can be initiated solely through direct recruitment via DNA, after which PRC2.1/PRC2.2 and PRC2/PRC1 functional interactions are required to achieve the full establishment of Polycomb binding through self- and mutual reinforcement.

**Figure 6.**
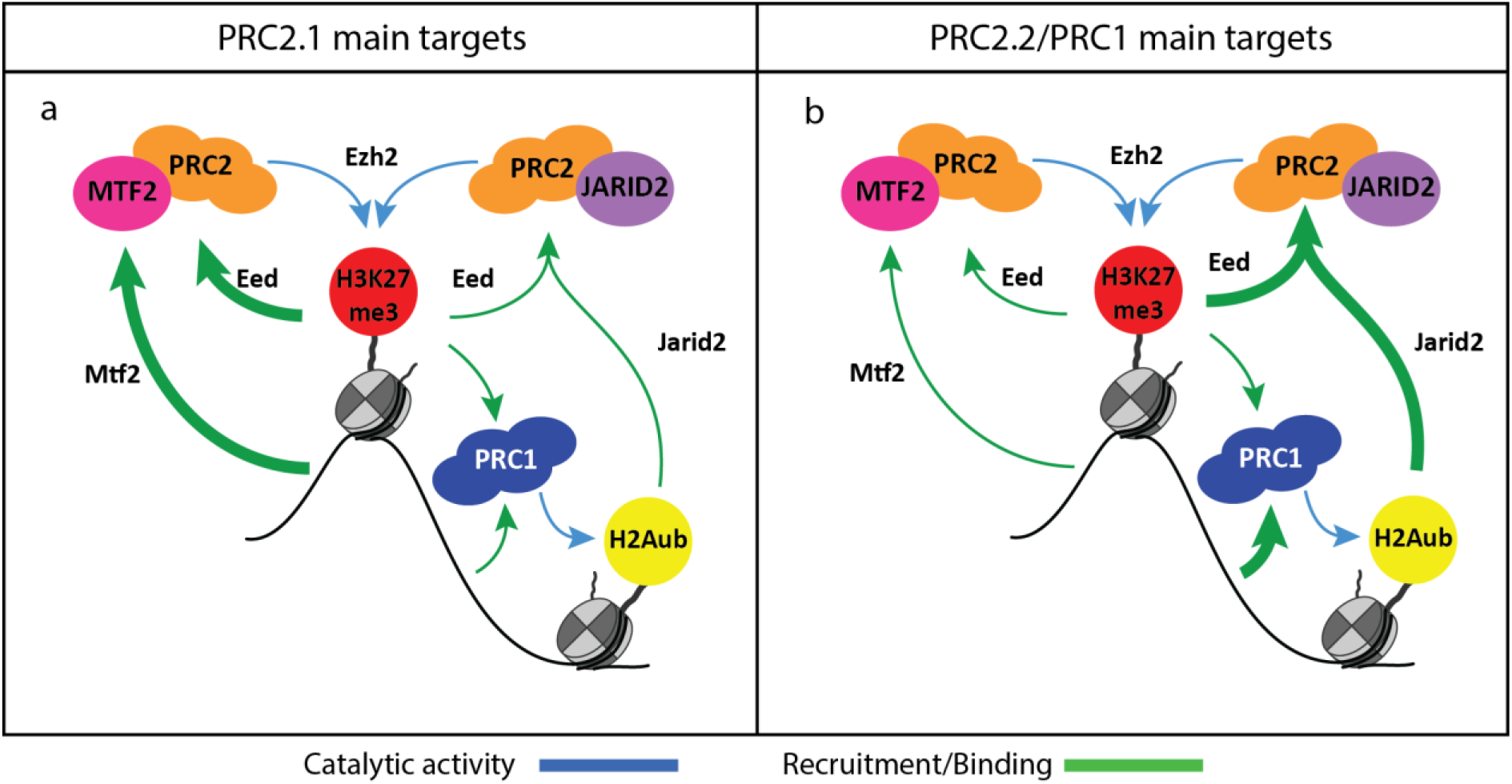
Model of PRC2 recruitment mechanisms and interactions. **a)** On PRC2.1 main targets (clusters 1-4) relatively little MTF2 binding is sufficient to kick start the EED positive feedback loop which heavily relies on JARID2. As primary recruitment is mediated to a large extent via MTF2, such a loop can still exist in the absence JARID2. In the absence of H3K27me3, an alternative route can take over that requires JARID2 binding to H2AK119ub. **b)** On PRC2.2/PRC1 targets (clusters 5-6), instead, Polycomb binding is initiated by PRC1 that, upon H2AK119ub deposition, is followed by JARID2-containing PRC2.2. These regions also see the presence of MTF2 in physiological conditions, but this is the result of indirect recruitment via the PRC2 core binding to PRC2.2-initiated H3K27me3 deposition.

## Acknowledgements

This work has been financially supported by the People Program (Marie Curie Actions) of the European Union’s Seventh Framework Program FP7 under grant agreement number 607142 (DevCom). G.v.M. is supported by the Oncode Institute, which is partly funded by the Dutch Cancer Society (KWF). H.M. is supported by the Netherlands Organisation for Scientific Research (NWO-VIDI 864.12.007). This work was carried out on the Dutch national e-infrastructure with the support of SURF Cooperative.

## Data availability

New and reanalysed published data are available via GEO (in submission), proteomics data via PRIDE (in submission). Normalized sequencing tracks can be browsed by loading the following track hub to the UCSC genome browser (XXX).

---

**Supplementary Figure 1.**
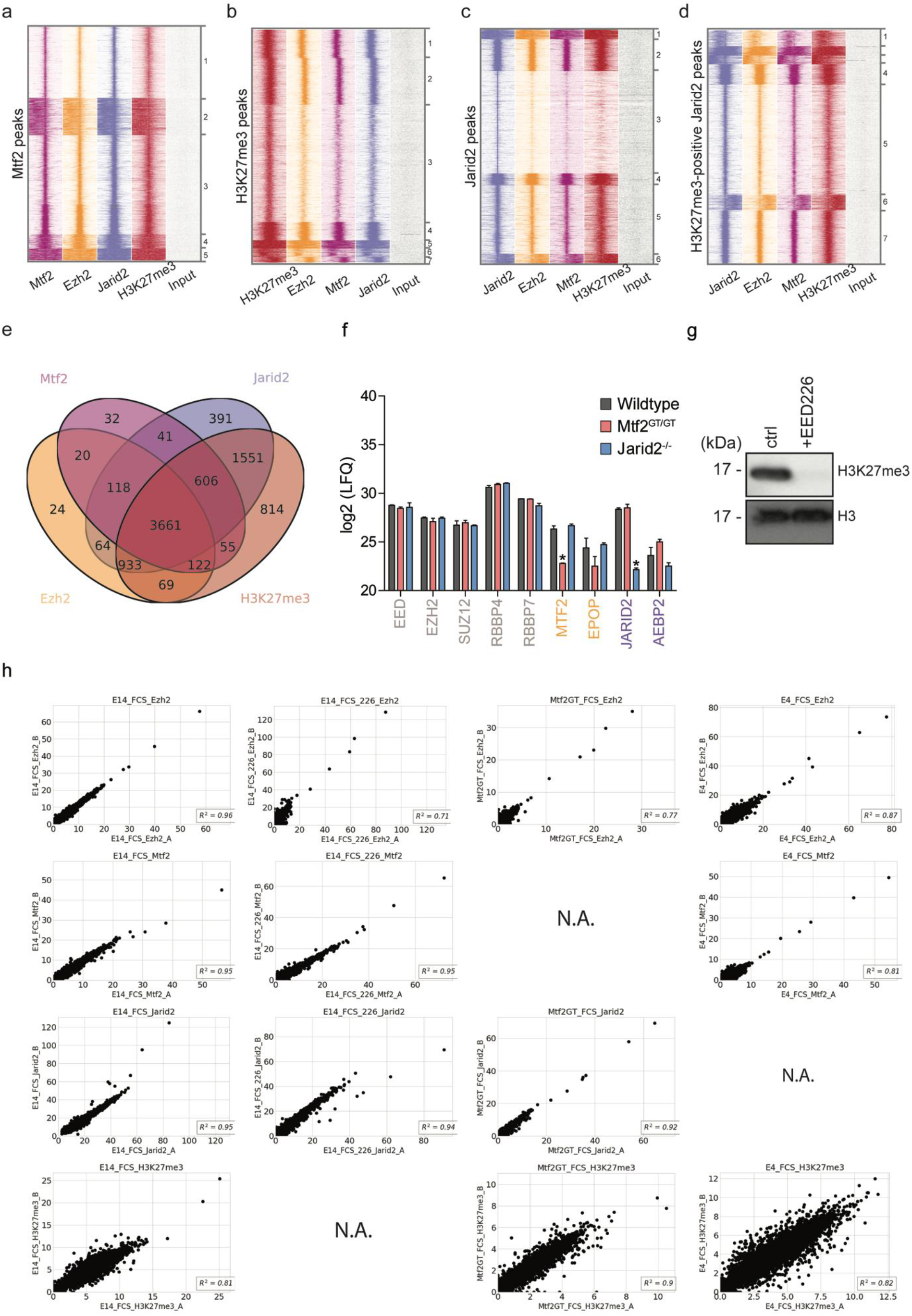
**a-d)** Heatmap of WT ChIP-seq signal on the indicated peak set. H3K27me3-negative JARID2 peaks were excluded from further analysis. **e)** Venn diagram showing the overlap of peaks called for the ChIP-Seq of each protein independently. **f)** Mass spectrometry quantification of PRC2 subunits in the different cell lines. Detection of JARID2 and MTF2 in the respective mutants (asterisks) is due to value imputation in Perseus. **g)** Western blot validation of EED226 depletion of H3K27me3, for the ChIP shown in Figs 1 and 2. **h)** Scatterplot of peak RPKM showing high reproducibility of ChIP replicates.

**Supplementary Figure 2.**
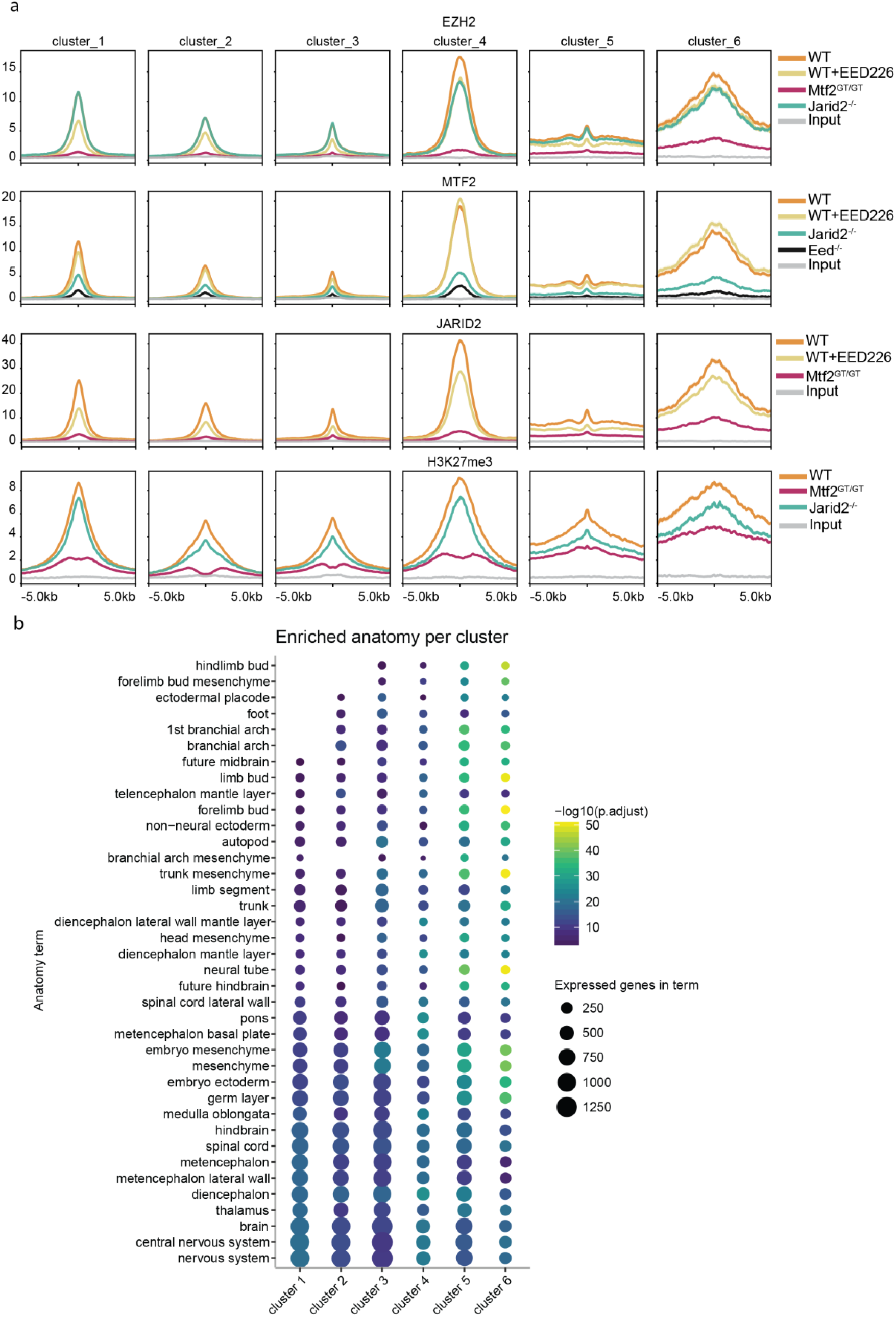
**a)** Average plot of the ChIP signal shown in Fig 2a, centred on called peaks. **b)** Enrichment of anatomical terms in the genes associated with peaks in the six clusters shown in Fig 2a. Enrichment over all genes.

**Supplementary Figure 3.**
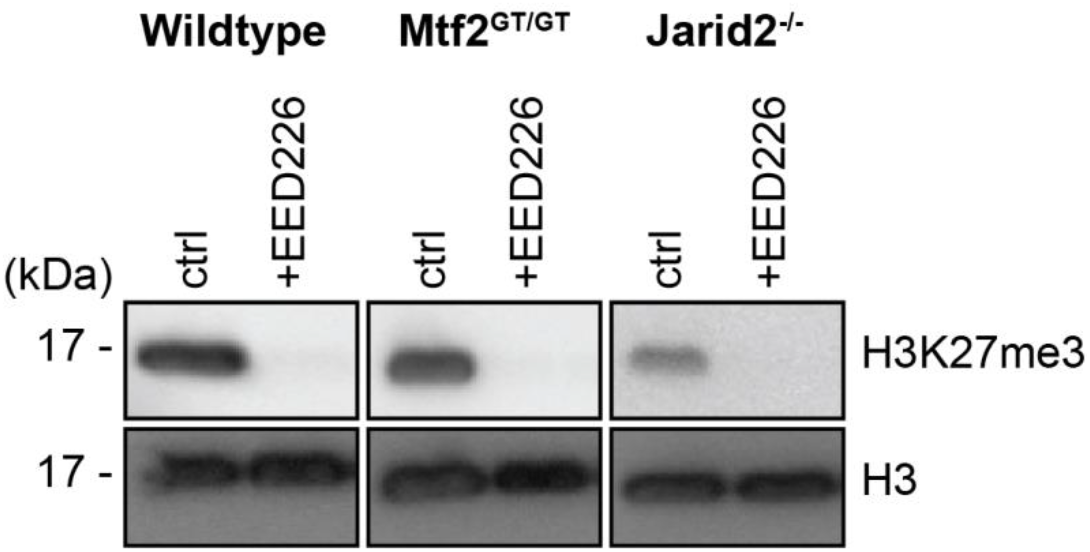
Western blot validation of EED226 depletion of H3K27me3 for the ChIP show in Figs 3 and 4

**Supplementary Figure 4.**
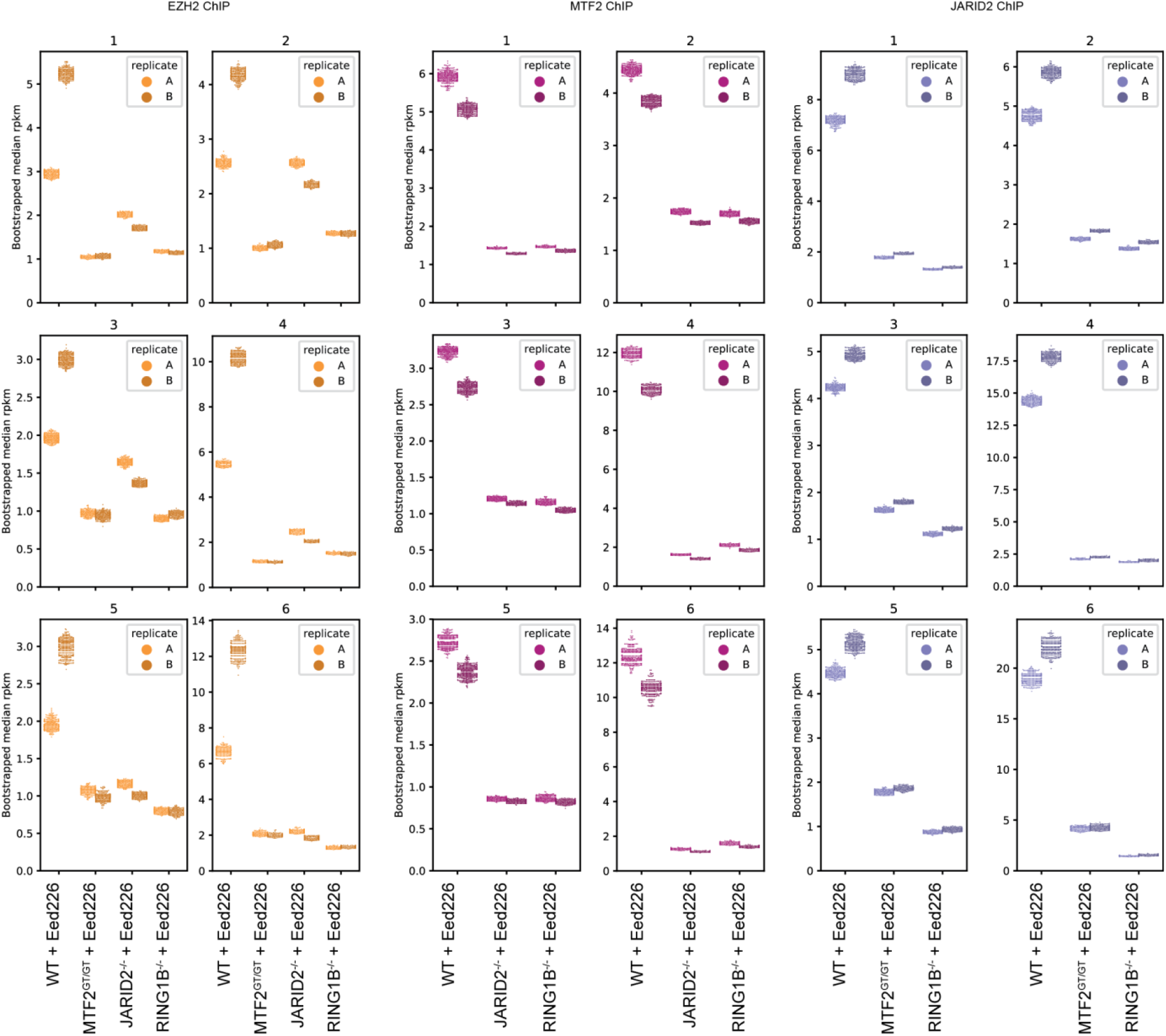
Bootstrapping-based RPKM quantification (methods) of the signal in Fig 4 a-d. Each coloured dot represent the median of one round of bootstrapping. Replicates are plotted independently.

**Supplementary Figure 5.**
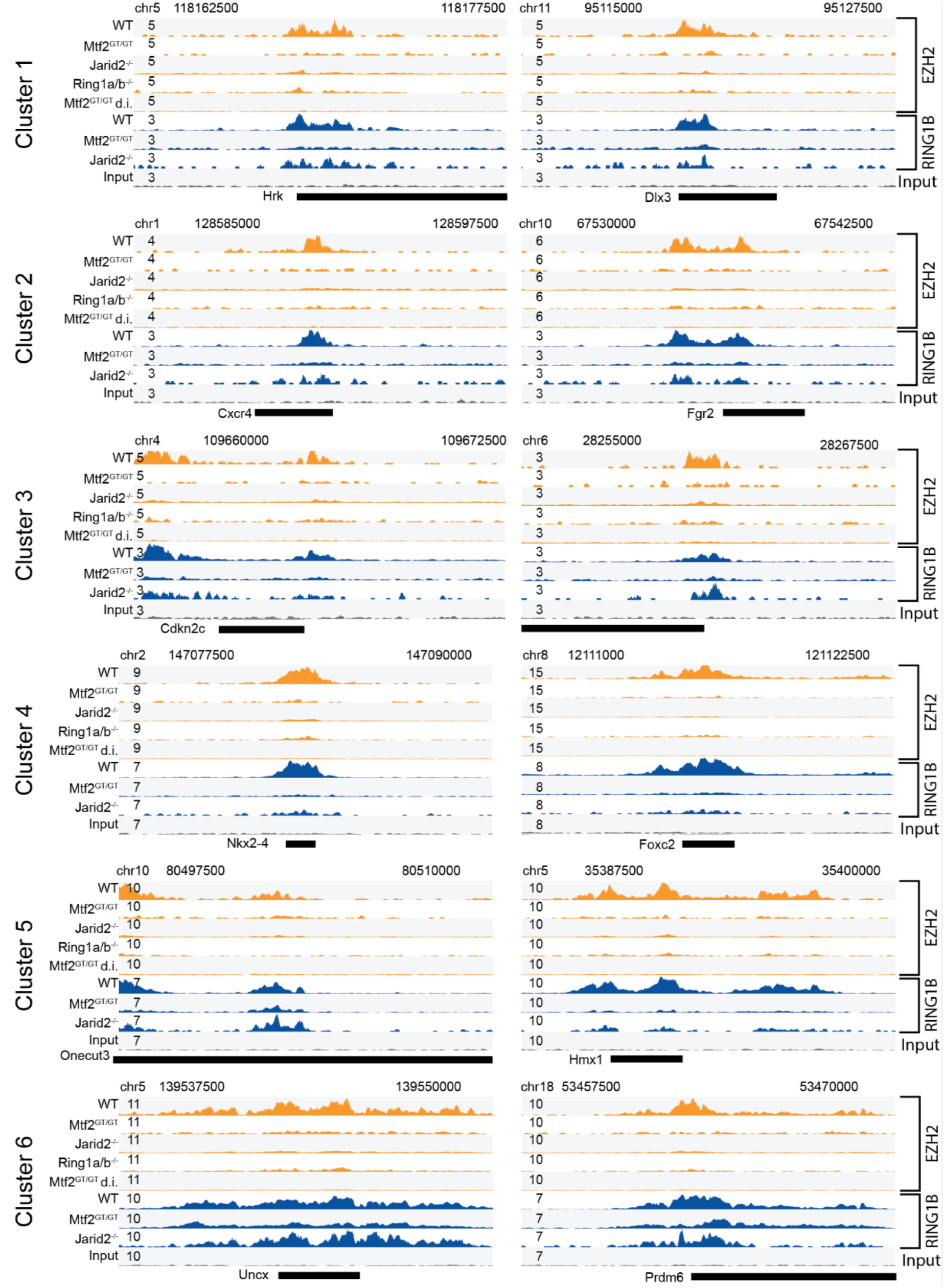
Examples loci of the data shown in Fig 4e-f and Fig5, two for each cluster.

